# Environmental DNA from archived leaves reveals widespread temporal turnover and biotic homogenization in forest arthropod communities

**DOI:** 10.1101/2022.04.27.489699

**Authors:** Henrik Krehenwinkel, Sven Weber, Rieke Broekmann, Anja Melcher, Julian Hans, Ruediger Wolf, Axel Hochkirch, Susan Rachel Kennedy, Jan Koschorreck, Sven Kuenzel, Christoph Müller, Rebecca Retzlaff, Diana Teubner, Sonja Schanzer, Roland Klein, Martin Paulus, Thomas Udelhoven, Michael Veith

**Affiliations:** Trier University, Trier, Germany; LMU Munich, Munich, Germany; German Federal Environment Agency, Berlin, Germany; Max Planck Institute for Evolutionary Biology, Ploen, Germany

## Abstract

A major limitation of current reports on insect declines is the lack of standardized, long-term, and taxonomically broad time series. Here, we demonstrate the utility of environmental DNA from archived leaf material to characterize plant-associated arthropod communities. We base our work on several multi-decadal leaf time series from tree canopies in four land use types, which were sampled as part of a long-term environmental monitoring program across Germany. Using these highly standardized and well-preserved samples, we analyze temporal changes in communities of several thousand arthropod species belonging to 23 orders using metabarcoding and quantitative PCR. Our data do not support widespread declines of *α*-diversity or genetic variation within sites. Instead, we find a gradual community turnover, which results in temporal and spatial biotic homogenization, across all land use types and all arthropod orders. Our results suggest that insect decline is more complex than mere *α*-diversity loss, but can be driven by *β*-diversity decay across space and time.

## Introduction

Dramatic declines of terrestrial insects have been reported in recent years, particularly in areas of intensified land use ^1–4^. However, some authors have urged caution in generalizing these results ^5–7^, suggesting that reported patterns of decline may be more localized than currently assumed or reflect long-term natural abundance fluctuations ^8,9^. Most studies on insect decline suffer from a lack of long-term time series data and are limited in geographic and taxonomic breadth, often using biomass as a proxy for diversity estimates ^1,10^. Hence, what is needed are methods and sample types that yield standardized long-term time series data for the diversity of arthropod communities across broad taxonomic and geographic scales ^11^.

In recent years, environmental DNA metabarcoding has offered a promising new approach to monitor biological communities ^12–14^. This includes terrestrial arthropods, whose eDNA can be recovered from various substrates, for example plant material ^15^. Here, we develop a DNA metabarcoding and quantitative PCR (qPCR) protocol to simultaneously recover diversity and relative DNA copy number of arthropod community DNA from powdered leaf samples. We then analyze 30-year time series data of arthropod communities from canopy leaf material of four tree species from 24 sites across Germany. These sites represent four land use types with different degrees of anthropogenic disturbance: urban parks, agricultural areas, timber forests, and national parks (Fig. 1). The samples were collected using a highly standardized protocol by the German Environmental Specimen Bank (ESB), a large biomonitoring effort for Germany’s ecosystems, and stored at below -150°C. By basing our analyses on DNA sequences, we can measure diversity from haplotype variation within species to taxonomic diversity of the whole community.

**Fig. 1:**
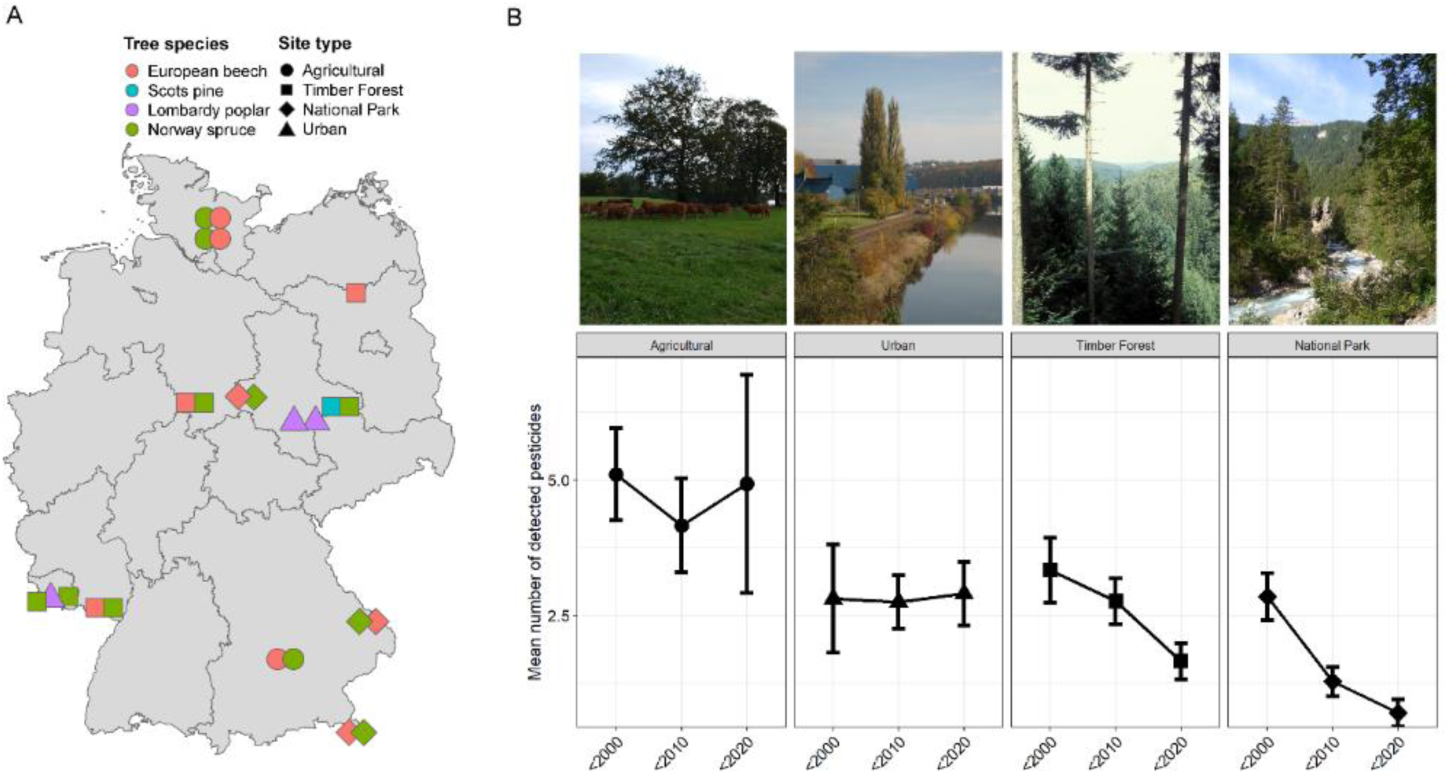
A) Sampling sites, land use types and the four tree species (European Beech *Fagus sylvatica*, Lombardy Poplar *Populus nigra* ‘Italica’, Norway Spruce *Picea abies*, and Scots Pine *Pinus sylvestris*) sampled by the ESB in Germany. B) Representative images of sampling sites for the four land use types and average number of detected pesticides across these land use types in three time periods (before 2000, 2000 – 2009, and 2010 – 2018). GC-MS/MS analysis ^23^ shows that the detected pesticide load distinguishes the different land use types, with agricultural sites continuously showing the highest number of pesticides.

Current studies on insect decline primarily focus on site-based assessments of *α*-diversity and biomass ^1,3^. Yet, these metrics alone are insufficient for characterizing ongoing biodiversity change ^16–18^. Significant temporal community change and declines can also occur at the scale of *β*- or γ-diversity, without affecting local richness. This may be driven by community turnover and spatial biotic homogenization ^19^. But diversity may also vary temporally. Fluctuating occurrences of transient species considerably increase diversity within single sites over time ^20^. The loss of such transient species in favor of taxa with a temporally stable occurrence results in an increasingly predictable community and hence a temporal diversity decline. Biotic turnover may occur gradually following changing environmental conditions, but can also occur abruptly, when ecosystems reach tipping points ^21^. In the latter case, a rapid and considerable biotic remodeling of the ecosystem may be found. The relevance of these spatial and temporal factors in insect decline remains largely elusive.

Here, we use our high-resolution data on arthropods from canopy leaf samples to test the hypotheses that **1)** *α*-diversity and biomass of canopy-associated arthropod communities have declined in the last 30 years ^1,3^, or **2)** community change has occurred in the form of turnover and possibly homogenization of communities across space and time ^17,22^. Last, we hypothesize that **3)** biodiversity declines will be particularly pronounced in areas of intensified land use.

## Results

### A standardized protocol to characterize plant-associated arthropod communities

We developed a standardized and robust protocol to reproducibly recover plant-associated arthropod communities from powdered leaf material. We controlled for effects of the amount of leaf-material per sample, rainfall before sampling, amount of leaf homogenate used for DNA extraction, extraction replication and primer choice on the recovered diversity (see Methods and Suppl. Figure 1 - 3 for details on standardization).

Using our optimized protocol, we analyzed 312 ESB leaf samples. We recovered 2,054 OTUs from our samples, with different tree species having significantly different OTU numbers on average (Fig. 2A, LMM, P < 0.05). Nevertheless, all tree species showed a balanced and relatively similar taxonomic composition at the order level (Fig. 2A, Suppl. Fig. 3E). We identified 23 orders, 218 families and 413 genera. The richest order was Diptera (600 OTUs in 48 families), followed by Hymenoptera (369 OTUs in 21 families), Acari (293 OTUs in 21 families), Lepidoptera (233 OTUs in 32 families), Hemiptera (152 OTUs in 19 families), Coleoptera (133 OTUs in 29 families), and Araneae (99 OTUs in 15 families). The recovered species assemblages were ecologically diverse, including herbivores, detritivores, predators, parasites and parasitoids (Suppl. Fig. 4A & 4B). Each tree species harbored a unique arthropod community (Fig. 2B, Suppl. Fig 3C & 3D), with typical monophagous taxa exclusively recovered from their respective host trees. The arthropod communities from different sites and land use types were also differentiated within tree species (Suppl. Fig. 5, PERMANOVA, P < 0.05). In addition to arthropod-host plant associations, we were able to detect interactions between arthropods. For example, abundances of the spruce gall midge *Piceacecis abietiperda* and its parasitoid, the chalcid wasp *Torymus* sp., were well correlated across all analyzed spruce sites (LM, P < 0.05). Both underwent coupled abundance cycles, with similar maxima every 6-8 years (Fig. 2C).

**Fig. 2:**
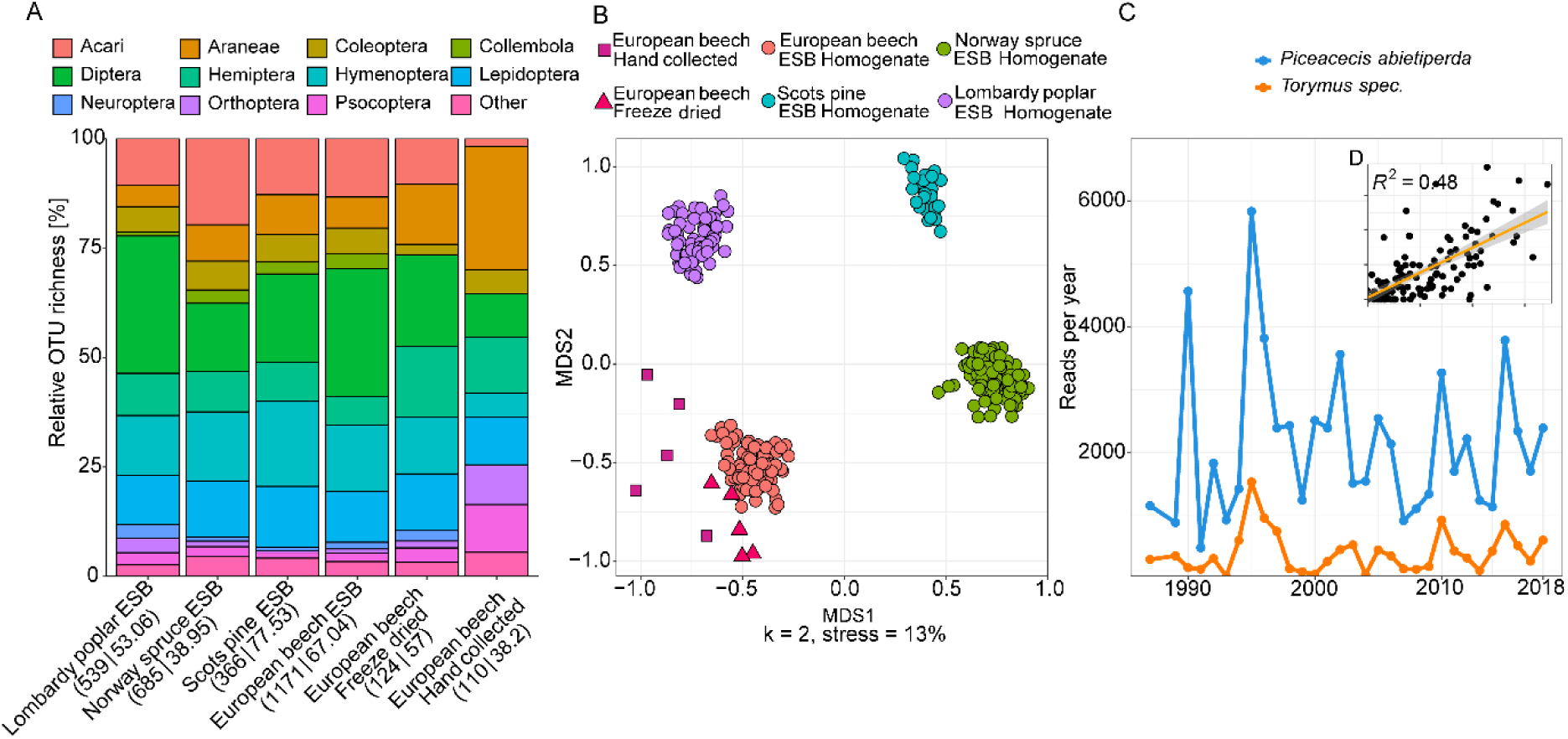
Recovery of diversity and interactions in canopy-associated arthropod communities from leaf material. A) Barplot showing recovered order composition of OTUs across the four tree species. In addition to ESB samples, results from freeze-dried leaves stored at room temperature for 6-8 years and hand-collected bulk insect samples from European Beech are shown. Orders amounting to less than 1 % of the total OTU number are merged as “Other”. Numbers below each barplot show the total number of arthropod OTUs followed by the mean OTU number per sample. B) NMDS plot showing tree-specific composition of arthropod communities for the same samples. C) Temporal changes in abundance of the spruce gall midge *Piceacecis abietiperda* and its parasitoid *Torymus* sp. between 1987 and 2018 in a spruce forest in the Saarland. Inset shows correlation of relative abundance between the two species across all ESB spruce samples.

The recovered community composition from ESB leaf homogenate samples was similar to hand-collected branch clipping samples (Fig. 2A+B). Branch clipping recovered a larger diversity of spiders, a taxon which is exclusively found on leaf surfaces. In contrast, about 25 % of the recovered taxa from ESB leaf powder likely inhabited the insides of leaves, e.g., gallers and miners (Suppl. Fig. 4B). Overall, arthropod DNA in leaf homogenates appears temporally very stable: Even freeze-dried leaf material that had been stored at room temperature for eight years yielded surprisingly similar arthropod communities to ESB samples (Fig 2A+B).

Besides analyzing diversity, we generated information on relative arthropod rDNA copy number in relation to the corresponding plant rDNA copy number by qPCR. Relative eDNA copy number should be a predictor for relative biomass. We designed a standardized qPCR assay based on lineage-specific blocking SNPs in the 18SrDNA gene (Fig. 3A). We tested two primer combinations with 3’-blocking SNPs in 1) only the forward or 2) both forward and reverse primer sequences. The primer combination with mismatches in both forward and reverse primers led to a near complete suppression of non-arthropod amplification in all tested samples (5.41 % vs. 44.49 % on average) and was hence chosen for the qPCR experiment (Fig. 3A+B). The qPCR assay showed a high efficiency (E_Plant_ = 94.77 %, E_Arthropod_ = 99.73 %) and accurately predicted changes in relative copy number of arthropod DNA on plant material, even when comparing taxonomically heterogeneous mock communities (Fig. 3C, Suppl. Fig. 6). Overall, we found a significant positive correlation of OTU richness and relative arthropod copy number in the ESB samples (LMM, P < 0.05; see Methods: “*Statistical analysis”*), supporting recent work suggesting a biomass-diversity relationship ^24^.

**Fig. 3:**
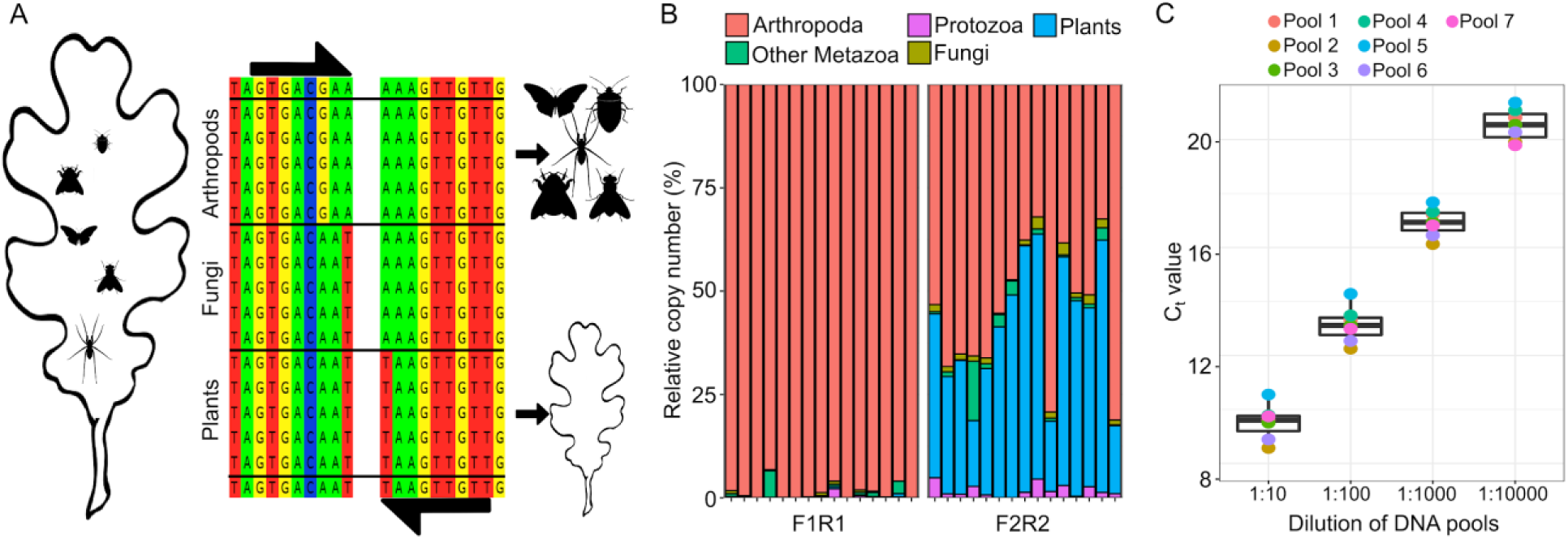
qPCR experiment to detect relative rDNA copy number of arthropods in plant homogenates. A) Schematic overview of the blocking approach to amplify homologous 18SrDNA fragments for either arthropod or plant DNA, based on lineage-specific priming mismatches. B) Effect of primer mismatches on the recovery of arthropod sequences. Barplots show recovered read proportion of different higher taxa from 15 ESB leaf samples. The left plot shows the effect of a diagnostic mismatch in forward and reverse primers, while the right plot shows the effect of only a forward primer mismatch. C) Boxplot showing CT values recovered from the seven mock communities of arthropod species from 13 different orders (see Suppl. Fig 6 for community composition) and across a 1:10000 dilution series. Separate CT values for each community are indicated by the dots.

### Temporal changes of diversity, copy number and species composition in canopy arthropod communities

Based on our time series data of archived ESB leaf samples, we tested the hypothesis that *α*-diversity (including intraspecific genetic diversity) and biomass (relative rDNA copy number) have undergone widespread temporal declines, particularly in areas of intensive land use. Our statistical analysis does not support previously reported widespread temporal *α*-diversity declines (Fig. 4A, Suppl. Fig. 3H & I, LMM, P > 0.05), even when different land use types are analyzed separately (Suppl. Fig. 7, LMM, P > 0.05). The temporal pattern of diversity was also largely independent of taxonomy: most orders did not show temporal trends when analyzed separately (Fig. 4C; Suppl. Fig. 8). Exceptions include a significant loss of lepidopteran diversity, which is primarily driven by OTU loss at urban sites, and an overall increasing diversity of mites (LMM, P < 0.05). The overall temporally stable diversity is also visible at separate sites; a diversity decline across all orders was observed at only a single site (Suppl. Fig. 9). Similar to *α*-diversity, we did not find widespread temporal declines of genetic diversity. Neither community-level zero radius OTU (zOTU) richness (which was well correlated to OTU richness, R^2^ = 0.89) nor within-OTU haplotype richness declined significantly over time (Fig. 4B; Suppl. Fig. 7B & 7C).

**Fig. 4:**
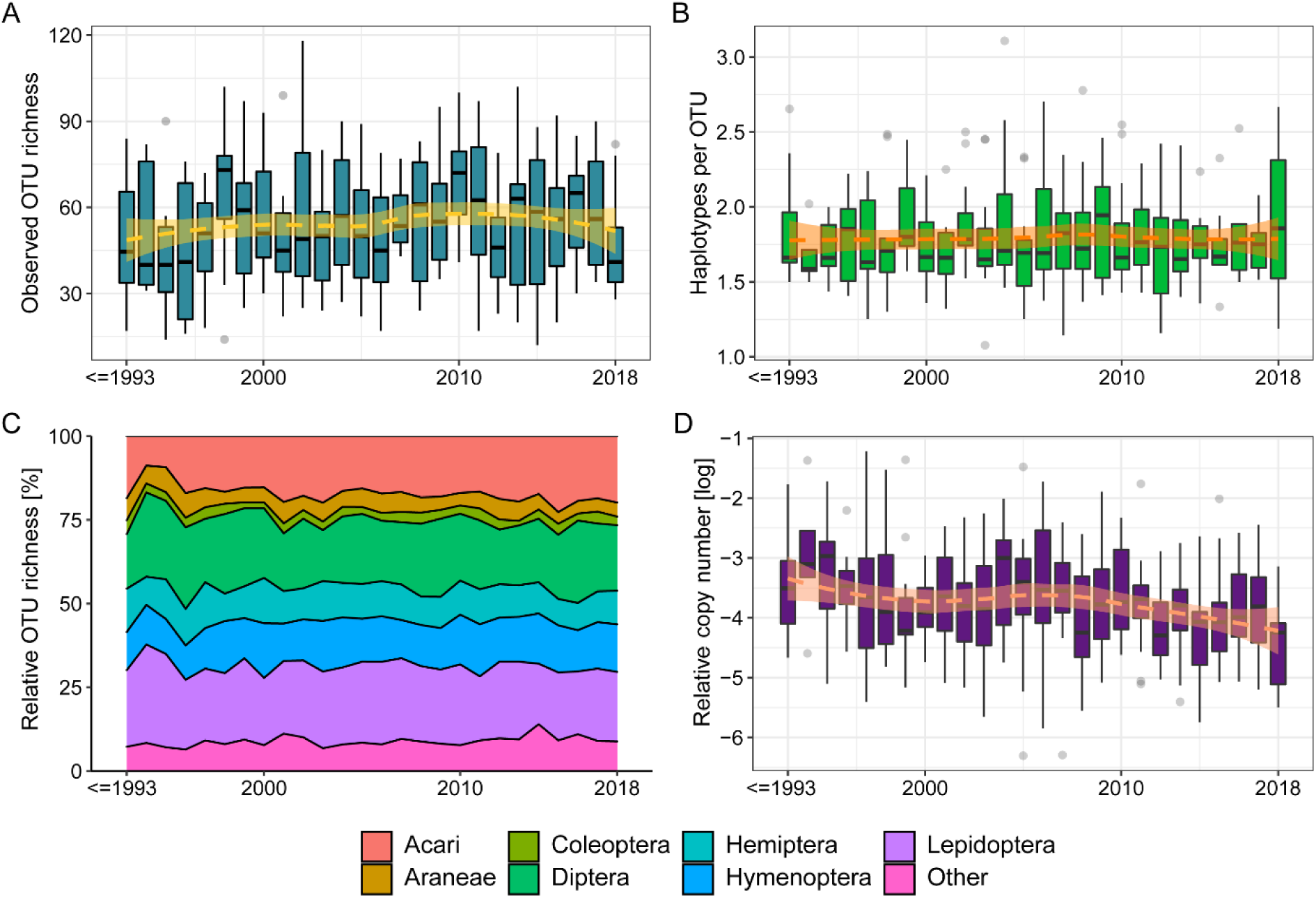
Temporal changes of diversity and copy number across all sampled sites. A) Arthropod OTU richness (representing *α*-diversity). B) Haplotype richness within OTUs (representing genetic variation). C) Relative OTU richness per order. D) Relative copy number of arthropod DNA (representing biomass).

In contrast to the stable *α*-diversity, relative copy number showed an overall decrease over time (Fig. 4D, LMM, P < 0.05), suggesting that arthropod biomass may indeed be declining in woodlands ^3^. The effect appears to be particularly driven by urban sites, coinciding with a loss of lepidopteran diversity (Suppl. Fig. 7-9). However, declines of arthropod DNA copy number are also visible in several agricultural and timber forest sites, particularly in the last 10 years of our time series (Suppl. Fig. 7 & 9).

We next explored temporal changes in abundance for 413 separate OTUs from a total of 19 sites. In line with our hypothesis 1), we predicted a majority of declining species. However, we found no significant difference between the average number of declining (6.94 %) and increasing (10.04 %) OTUs (t-test, P > 0.05, Fig. 5A). Declines and increases in OTU read abundance were also independent of arthropod order and land use type (Fig. 5A+B, Fisher’s exact test P > 0.05). The observed replacement of about 15 % of OTUs within sites translates into a significant temporal change of taxonomic *β*-diversity (Fig. 5C, Suppl. Fig. 3J). We found a strong positive correlation of temporal distance and Jaccard dissimilarity for most sites (PERMANOVA, P < 0.05). Thus, species are continuously replaced in all land use types (Suppl. Fig. 7E).

**Fig. 5:**
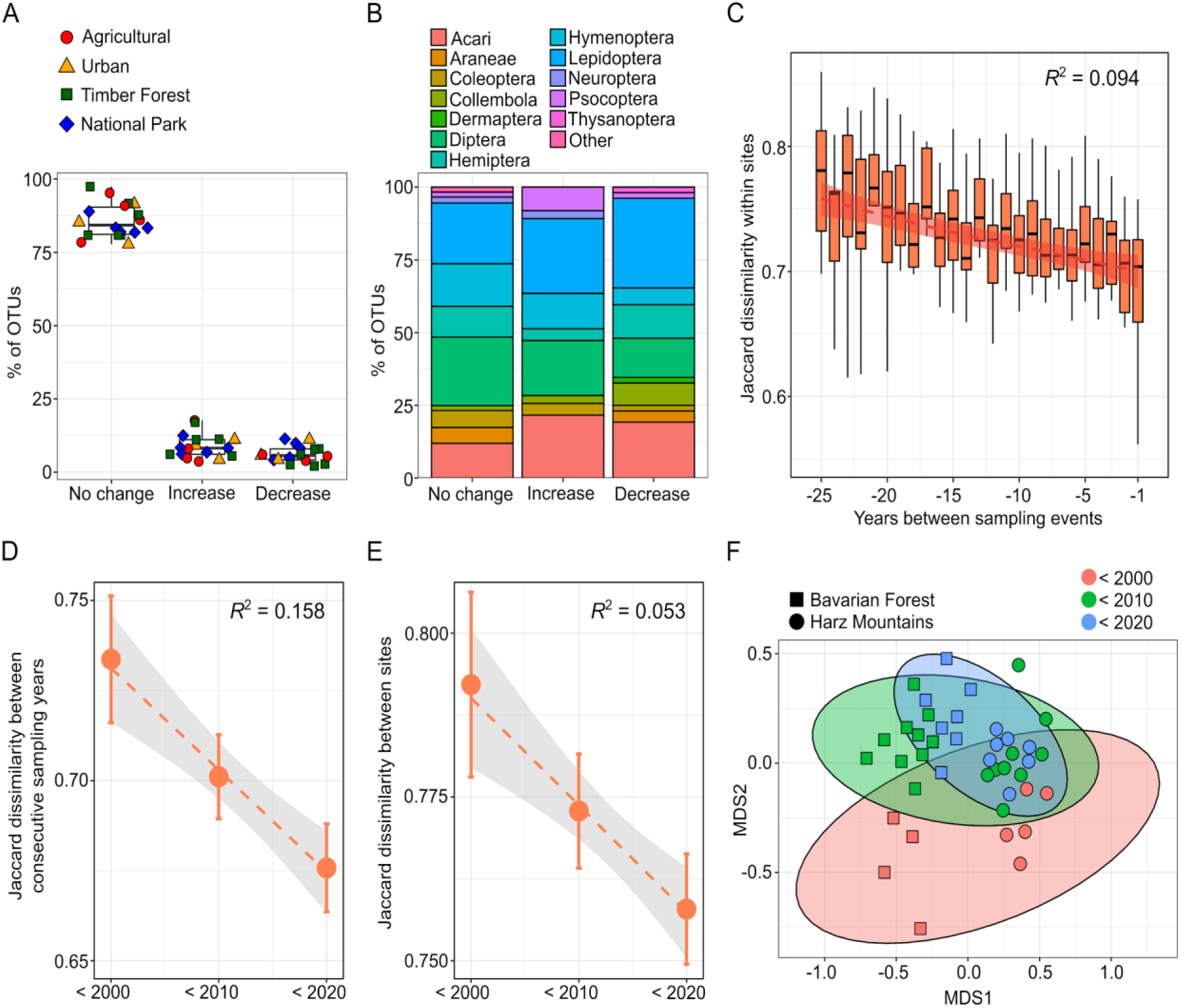
Temporal changes of species composition and *β*-diversity within and between sampling sites. A) Boxplots of the proportion of 413 separate OTUs that significantly increased, decreased, or did not show temporal abundance changes. The colored symbols overlaying the boxplots represent land use type for each site. B) Stacked barplot showing the recovered order composition of the same three categories of OTUs. Orders amounting to less than 1 % of the total OTU number are merged as “Other.” C) Within-site Jaccard dissimilarity as a function of number of years between sampling events, showing a pronounced taxonomic turnover. D) Jaccard dissimilarity between consecutive sampling years in each decade of sampling, calculated within sites. We find a loss of temporal *β*-diversity over time. Points indicate means and error bars show 95% confidence interval. E) Jaccard dissimilarity between sites. We find a loss of average between-site *β*-diversity, i.e. spatial homogenization. F) NMDS plot showing community dissimilarity within and between Bavarian Forest and Harz Mountains national parks over three decades.

While the community turnover did not affect local *α*-diversity, we still observed associated losses of overall diversity. The first noteworthy pattern concerns a loss of temporal *β*-diversity within sites. *β*-diversity between consecutive sampling years dropped significantly in many sites, particularly in beech forests (Fig. 5D). Thus, diversity within sites is increasingly homogenized over time. We also found a significant decrease of *β*-diversity between sites for beech forests (Fig. 5E). Our data suggest a loss of site-specific species and a gain of more widespread generalists, irrespective of land use. This pattern also emerges at the level of individual OTUs. Several novel colonizers spread rapidly in woodlands and showed similar abundance trends across various sites in parallel (Suppl. Fig. 10). The spatial and temporal change of *β*-diversity is illustrated by an NMDS plot of arthropod communities from two beech forests in National Parks, the Harz and the Bavarian Forest (Fig. 5F). While the two sites are well separated by the first NMDS axis, the second axis shows a pronounced temporal turnover of communities. In the past decades, this turnover has led to temporally more predictable communities within sites and increasingly similar communities between the two national parks.

## Discussion

Here we show that DNA from archived leaf material provides a robust source of data to reconstruct temporal community change across the arthropod tree of life. Leaf samples should also cover a broad phenological window: Adults of many insect species are only active during a short time period of the year, but their larvae spend the whole year on their host plant ^25^. As sampled leaves make up the habitat and often food source of arthropods, it is also possible to infer the exposure of arthropod communities to chemical pollution by analyzing chemicals in the leaf material. This is of critical importance, as pesticide use has often been invoked as a driver of insect decline ^26,27^. The high temporal stability of arthropod DNA in 8-year-old dried leaf samples also highlights the utility of other plant archives, e.g., herbaria, for arthropod eDNA.

We here analyze a unique leaf archive and provide unprecedented insights into arthropod community change in the tree canopy, an ecosystem known for its high and often cryptic diversity ^15^. Our results do not confirm the hypothesis of widespread losses of *α*-diversity. Initial reports of insect declines originate mainly from grassland ecosystems that have undergone massive changes in land use ^1^. Central European canopy communities may be less affected by such change. Interestingly, the only sites that showed declines of richness were agricultural and urban sites, suggesting that land use may at least locally affect neighboring canopy communities ^28^. However, we detected DNA copy number declines in all land use types despite a temporally stable *α*-diversity, suggesting that biomass may indeed be in decline. Alternatively, the dropping copy number may also reflect a taxonomic turnover, of species with different eDNA shedding rates.

Instead of widespread losses of *α*-diversity, we indeed found pronounced taxonomic turnover in nearly all communities ^22^, supporting our second hypothesis. While resident species are continuously lost, they are mostly replaced by novel colonizers. This turnover can result in biotic homogenization across space and time. Less interannual variation of the occurrence of taxa within sites and reduced spatial variation of their occurrence between sites both cause a decline in overall *β*-diversity. Biotic homogenization is often associated with intensification of land use ^19,29^ and landscape simplification ^30^. However, the pattern we observed affected all land use types equally. The universality of these changes suggests that neither site- nor taxon-specific factors are responsible. Possible explanations include factors that act at a larger scale, such as climate change-induced range shifts ^16^, nitrogen deposition ^31^ and the introduction of invasive species ^32,33^. Given that our leaf samples recover a fairly broad phenological window, the alternative explanation that the observed community-wide turnover pattern may have resulted from shifting phenologies ^34^ is unlikely. The gradual replacement of species also suggest that we are not yet observing ecosystems reaching tipping points ^21^.

In summary, our work shows the great importance of standardized time series data to accurately reveal biodiversity change in the Anthropocene across space and time, beyond the decline of *α*-diversity and biomass ^22^. Taxonomic replacement and biotic homogenization, even in seemingly pristine habitats such as national parks, signify an important and hitherto insufficiently recognized facet of the current insect crisis.

## Materials and Methods

### Samples and metadata used in this study

#### Tree samples of the German Environmental Specimen Bank – Standardized time series samples stored at ultra-low temperatures

We used a total of 312 leaf samples of four common German tree species: the European Beech *Fagus sylvatica* (98 samples), the Lombardy Poplar *Populus nigra “italica’* (65 samples), the Norway Spruce *Picea abies* (123 samples), and the Scots Pine *Pinus sylvestris* (26 samples). The samples have been collected annually or biannually by the German Environmental Specimen Bank (ESB) since the 1980s and serve as indicators for aerial pollutants ^35^. A total of 24 sampling sites were included, covering sampling periods of up to 31 years and representing four land use types of varying degrees of anthropogenic disturbance (Fig. 1). These include natural climax forest ecosystems in core zones of national parks (6 sites, National Park), forests commercially used for timber (9 sites, Timber Forest), tree stands in close proximity to agricultural fields (6 sites, Agricultural), and trees in urban parks (3 sites, Urban). The sites were initially chosen to represent their land use type permanently for long term monitoring, and the corresponding land use categories have mostly remained temporally stable.

The samples have been collected and processed according to a highly standardized protocol at the same time every year and using sterile equipment ^36–38^. A defined amount of leaf material (> 1.100 g) is collected from a defined number of trees (>10) from each site and a defined number of branches from each tree. This amount translates to several thousand leaves from each site, which should suffice to saturate the recovered arthropod diversity. The sampled leaves are intended to represent the exact natural state of the tree. They are not washed or altered in any way before processing, and eDNA traces, and small arthropods on the leaves’ surfaces, as well as from galls and leaf mines, are included in the sample. Each sample is stored on liquid nitrogen immediately after collection and ground to a powder with an average diameter of 200 µm using a cryomill. The resulting homogenates are stored for long term on liquid nitrogen ^39,40^. The cold chain is not interrupted after collection and during processing, ensuring optimal preservation of nucleic acids in the samples. The homogenization of the sample also guarantees a thorough mixing, resulting in equal distribution of environmental chemicals and eDNA in the sample. Previous work suggest that very small subsamples of the homogenate suffice to detect even trace amounts of environmental chemicals in the sample ^35^. eDNA from leaf surfaces may be affected by weather conditions before sampling, e.g., washed away by heavy rain or damaged by strong UV exposure ^41^. However, leaves are only collected dry by the ESB, i.e., not immediately after rain. The date of the most recent rainfall is noted for each ESB sampling event, allowing us to explore the effects of recent weather conditions on the recovered arthropod communities.

#### The utility of plant material stored at room temperature to recover arthropod DNA

ESB samples are stored under optimal conditions for nucleic acid preservation. By contrast, most archived leaf samples are stored at room temperature, e.g. dried leaf material in herbaria. To test the general suitability of archived leaf material for arthropod community analysis, we included 25 additional beech leaf samples, each consisting of 100 leaves from a total of five sites. The samples were freeze-dried and then ground to a fine powder by bead beating. The resulting powder was comparable to our ESB homogenates, but unlike them, it was stored at room temperature for 6-8 years.

#### Hand-collected branch clipping samples to explore the accuracy of leaf-derived arthropod DNA

To evaluate the performance of our leaf DNA based protocol in comparison to commonly used sampling methods, we generated a branch clipping dataset from five beech stands close to Trier University. Branch clipping is a widely used method to collect arthropods residing on leaf surfaces in trees ^42^ and thus the best comparable traditional methodology to our protocol. We sampled five trees per site and collected ten branches of about 40 cm length from each tree. The branches were clipped off, stored in plastic bags and then brought to the laboratory. Here, arthropods were manually collected from each sample. All collected arthropod specimens were pooled by tree and stored for later DNA extraction in 99 % ethanol.

#### Climate data

We downloaded monthly climate data for all study sites from the German Climate Center distributed as a raster dataset interpolated from the surrounding weather stations by the German Meteorological Service (Deutscher Wetterdienst – DWD). We collected data for average annual temperature and rainfall as well as summer and winter temperature and rainfall.

#### Measurement of pesticide content from archived leaf material

The ESB sampling was historically set up as a tool for pollution assessment ^35^, and the samples are therefore stored to preserve any possible pollutant. Because these leaves serve as a habitat for the associated arthropods, such well-preserved material allows us to explore levels of chemical pollution occurring directly within the arthropods’ environment. Our samples were screened for pesticides and persistent organic pollutants with a modified QuEChERS approach ^23^. 2.0 g of sample material were extracted with acetonitrile (10 mL) and ultrapure water (10 mL), followed by a salting-out step using magnesium sulfate, sodium chloride and a citrate buffer (6.5 g; 8:2:3 (w/w)). After a dispersive solid-phase extraction (dSPE) cleanup step with magnesium sulfate, PSA and GCB (182.5 mg; 300:50:15 (w/w)), the supernatant was analyzed with a sensitive gas chromatography-tandem mass spectrometry (GC-MS/MS) instrument. All samples were analyzed for 208 GC-amenable compounds of different pesticide and pollutant classes, including pyrethroids, organochlorine and organophosphate pesticides and polychlorinated biphenyls.

### Molecular processing

#### DNA isolation

We developed a highly standardized protocol for the analysis of leaf-associated arthropod community DNA. We optimized various protocol steps to ensure the reliability and reproducibility of our data. We first explored the effect of DNA extraction on recovered diversity. As mentioned above, the cryo-homogenization of ESB samples ensures a very homogeneous distribution of even trace amounts of chemicals in the sample. This should also hold true for DNA. Hence small subsamples of large homogenate samples should suffice for analysis. To test this hypothesis, we first performed a weight series extraction from 50, 100, 200, 400, 800 and 1600 mg of homogenate with several replicates for each weight. Additional extraction replicates of 16 beech samples at 200 mg were also included. This analysis confirms the pronounced homogenization of the samples, with 200 mg of homogenate sufficing to accurately recover *α*- and *β*-diversity (Suppl. Fig. 1A & 1B).

DNA was extracted from all ESB and freeze-dried samples, using the Puregene Tissue Kit according to the manufacturer’s protocols (Qiagen, Hilden, Germany). All samples were processed under a clean bench and kept over liquid nitrogen during processing to prevent thawing. Samples were transferred using a 1000 µL pipette with cut off tips. The resulting wide bore tips were used to drill out cores of defined sizes from the leaf powder. To remove undesired coprecipitates, we performed another round of purification for each sample, using the Puregene kit following the manufacturer’s protocol. The hand-collected arthropod specimens from our branch clipping were pulverized in a Qiagen Tissuelyzer at 200 hz for 2 min using 5 mm stainless steel beads, and DNA was extracted from the pulverized samples using the Puregene kit as described in de Kerdrel et al. ^43^. Branch clipping and leaf samples were processed in separate batches and using separate reagents to avoid possible carryover between sample types.

#### Primer choice, PCR amplification and sequencing

As the standard DNA barcode marker for arthropods ^44^, the mitochondrial COI gene offers the best taxonomic identification of German arthropod species. We thus selected COI for our metabarcoding analysis. We tested several primer pairs to optimize recovery of arthropod DNA from the leaf homogenates ^45–47^. Leaf homogenates are dominated by plant DNA, with arthropod eDNA only present in trace amounts. The majority of commonly used arthropod metabarcoding primers are very degenerate and will readily amplify mitochondrial COI of plants. Thus, for such degenerate primers, the vast majority of recovered reads will belong to the plant. We found an ideal tradeoff between suppressing plant amplification while still recovering a taxonomically broad arthropod community in the primer pair ZBJ-ArtF1c/ZBJ-ArtR2c ^48^. While this primer pair is known to have taxonomic biases for certain arthropod groups ^49^, it is still widely used as an efficient and reliable marker for community analysis ^14,50^. Recently, we designed a novel and highly degenerate primer pair by modifying two degenerate metabarcoding primers ^45,46^, which allows the suppression of plant amplification (*NoPlantF_270/mICOIintR_W*; See Suppl Table 1). To ensure the reproducibility of the diversity patterns recovered from our original ZBJ-ArtF1c/ZBJ-ArtR2c dataset, we additionally processed eleven complete ESB time series (174 samples) for this novel primer pair and compared results for species composition, *α*- and *β*-diversity. This analysis supports very similar patterns for temporal species abundance trends, as well as *α*- or *β*-diversity for both primer pairs (Suppl. Fig. 3).

All PCRs were run with 1 µL of DNA in 10 µL volumes, using the Qiagen Multiplex PCR kit according to the manufacturer’s protocol and with 35 cycles and an annealing temperature of 46 °C. A subsequent indexing PCR of 5 cycles at an annealing temperature of 55 °C served to attach sequencing adapters to each sample (using the layout described in Lange et al. ^51^). As technical replicates, all PCRs were run in duplicate. DNA extraction and PCR replicates showed well correlated and reproducible OTU composition (R^2^_extract1vs2_ = 0.90, R^2^_PCR1vs2_ = 0.97, LM P< 0.05), as well as *α*-diversity patterns (R^2^_extract1vs2_ = 0.93, R^2^_PCR1vs2_ = 0.90, LM P< 0.05). PCR- and extraction replicates also recovered a significantly lower *β*-diversity than within- and between-site comparisons (*β*_PCR1vs2_ = 0.14; *β*_extract1vs2_ = 0.19; *β*_within_site_ = 0.62; Pairwise Wilcoxon Test, P < 0.05) (Suppl. Fig. 1). The final libraries were quantified on a 1.5 % agarose gel and pooled in approximately equal abundances based on gel band intensity. The final pooled sample was cleaned using 1X Ampure Beads XP (Beckmann-Coulter, Brea, CA, USA) and then sequenced on an Illumina MiSeq (Illumina, San Diego, CA, USA) using several V2 kits with 300 cycles at the Max Planck Institute for Evolutionary Biology in Plön, Germany. Negative control PCRs and blank extraction PCRs were run alongside all experiments and sequenced as well. Branch clipping samples were amplified and sequenced separately using the above protocol.

#### Sequence processing

Demultiplexed reads were merged using PEAR ^52^ with a minimum overlap of 50 and a minimum quality of 20. The merged reads were then quality filtered for a minimum of 90 % of bases > Q30 and transformed to fasta files using FastX Toolkit ^53^. Primer sequences were trimmed off using *sed* in UNIX and the reads dereplicated using USEARCH ^54^. The dereplicated sequences were clustered into zero radius OTUs (hereafter zOTUs) using the *unoise3* command ^55^ and 3 % radius OTUs using the *cluster_otus* command in USEARCH. Chimeras were removed *de novo* during OTU clustering. All resulting sequences were translated in MEGA ^56^ and only those with intact reading frames were retained. To assign taxonomic identity to the zOTU sequences, we used BLASTn ^57^ against the complete NCBI nucleotide database (downloaded February 2021) and kept the top 10 hits. Sequences were identified to the lowest possible taxonomic level, with a minimum of 98 % similarity to classify them as species. All non-arthropod sequences were removed. We then built Maximum Likelihood phylogenies from alignments of the zOTU sequences for all recovered arthropod orders separately using RaxML ^58^. These phylogenies were used to perform another clustering analysis using ptp ^59^ to generate OTUs from the data. The ptp clustering came closest to the actual species assignments of zOTUs by BLAST, while 3 % radius OTUs tended to oversplit species. Moreover, the recovered diversity values for zOTUs and ptp-based OTUs were well correlated (Suppl. Fig. 1, R^2^ = 0.90). We thus proceeded to use ptp-based OTUs (hereafter referred to as OTUs) for subsequent analysis on taxonomic diversity. zOTUs represent individual haplotypes in the dataset and were thus used as an indicator of genetic diversity. Using the taxonomic assignments, we estimated which taxonomic groups were particularly well represented in our data. Each tree species likely harbors a unique arthropod community with numerous monophagous species, a majority of which should be recovered by a broadly applicable molecular method. Where possible, we performed a finer scale ecological assessment for the recovered taxa, classifying them by trophic ecology and expected position on the outside or inside of the leaf. For example, mining taxa would likely be recovered from the inside of the leaf, while other taxa likely reside on the leaf’s surface.

#### Detection of relative arthropod DNA copy number using quantitative PCR

Initial reports on insect decline were entirely based on biomass ^1^. Biomass, however, does not necessarily predict diversity ^60^. We therefore aimed to generate information not only on diversity, but also on relative biomass of arthropods in tree canopies. Previous eDNA studies show that DNA copy number is correlated with the biomass of a target taxon ^56^, making qPCR a possible approach for biomass estimation. We developed a qPCR protocol to detect relative abundance of arthropod DNA copy number in leaf samples, using the plant DNA copy number as an internal reference for quantification. We used the nuclear 18SrRNA gene (hereafter 18S). Although 18S can show interspecific copy number variation, it provides relatively good approximations of actual taxon abundances in amplicon assays ^61,62^. Primer pairs targeting plants and arthropods were designed to meet the following criteria: 1) Identical PCR fragments should be amplified for plants and arthropods so that PCR for both taxa will perform similarly. 2) The arthropod-specific primer should not amplify plants, and vice versa. 3) Fungi should be excluded from amplification, as DNA of fungal endophytes is probably at least as abundant in leaf samples as arthropod eDNA. 4) The primers should target conserved regions in order to amplify a broad spectrum of plants or arthropods. We used diagnostic SNPs at each primer’s 3’-end to achieve the lineage specificity ^63^.

Two possible qPCR primer pairs were designed, one targeting a 172 bp and the other a 176 bp fragment of 18S. The first primer pair contained a 3’-AA-mismatch discriminating arthropods from plants and fungi in the forward primer and a 3’-TT-mismatch discriminating plants from arthropods in the reverse primer, while the second pair had the same 3’-AA-mismatch in the forward primer but no reverse primer mismatch (Fig. 3A). To test the lineage specificity of both primer pairs, we performed an amplicon sequencing experiment with the arthropod-specific primers. Four samples from each of the four tree species were amplified with both primer pairs, then indexed, pooled, sequenced and processed as described above. All reads were clustered into 3 % radius OTUs. Taxonomy was assigned to the OTUs, and an OTU table was built, as described above for the ESB sample metabarcoding experiment. The proportion of arthropod reads was then estimated for each sample and primer pair. The first primer combination (3’-mismatch in both forward and reverse primers) led to a near complete suppression of plant and fungal amplification in all tested samples and was therefore used for the qPCR (Fig. 3A & 3B). Of this primer pair, two different versions were ordered: one arthropod-specific and the other plant-specific, both amplifying an identical stretch of 18S.

To account for the low quantity of arthropod DNA in relation to plant DNA, we designed a nested qPCR assay, with higher sensitivity for low DNA copy number ^64^. The sample was first amplified in a regular PCR with 15 cycles using the Qiagen Multiplex PCR kit. Two separate PCRs were run: one using the arthropod primers with an undiluted DNA extract as template, and the other using the plant primers with a 1:100 dilution of the DNA extract. The primers included a 33 bp forward and 34 bp reverse tail, based on Illumina TruSeq libraries, which were complementary to sequences in the qPCR primers. After being cleaned of residual primers with 1X AMPure beads XP, the products of the first PCR were used as template in the qPCR. qPCR was run with the Power SYBR Green Mastermix (Fisher Scientific, Waltham, MA, USA) on an ABI StepOnePlus Real-Time PCR System (Applied Biosystems, Foster City, CA, USA) according to the manufacturer’s protocol, using 35 cycles and an annealing temperature of 55 °C. All reactions were run in triplicate and the average of the three CT values used for analysis. The qPCR efficiency was estimated using a 10,000-fold dilution series. To estimate the accuracy of relative arthropod DNA copy detection across diverse arthropod communities, we also performed a spike-in assay, in which a serial dilution series (10,000-fold) of arthropod mock community DNA was added to a leaf extract and analyzed using qPCR. Seven mock communities were prepared, each containing varying amounts of DNA from 13 arthropod species representing 13 different orders (Suppl. Fig 8). The relative copy number of arthropod DNA in relation to plant DNA was estimated using the Delta CT Method ^65^. The optimized qPCR protocol was then used to quantify the relative DNA copy number of arthropods in all 312 ESB leaf samples.

#### Statistical analysis

Using USEARCH, an OTU table was built including all samples with the taxonomically annotated zOTU sequences as reference. Based on the negative control samples, we removed all entries in the OTU table with fewer than 3 reads. Using rarefaction analysis in vegan ^66^ in R (v 4.1.0) ^67^, we explored saturation of diversity. Based on this analysis, 5000 reads were randomly sampled for each of the two PCR duplicates using GUniFrac ^68^, and the duplicates were merged into a final sample of 10,000 reads after they were checked for reproducible patters of species composition, *α*- and *β*-diversity. Taxonomic *α*- and *β*-diversity were calculated in vegan in R. Our dataset contained different sources of DNA, e.g., eDNA and whole bodies, from different arthropod groups. Quantitative biodiversity assessments at the community level were thus likely biased. We therefore refrained from quantitative assessments of biodiversity and limited our assessments of *α*-diversity to richness and *β*-diversity to binary dissimilarity. As mentioned above, zOTUs represent individual haplotypes and thus genetic variation within species. Using the zOTU data, we calculated the haplotype (zOTU) richness within each individual OTU as a complementary measure for genetic variation. A decline in biodiversity could manifest itself in an overall loss of species, which should be detectable at the OTU level. Alternatively, biodiversity decline could initially only affect genetic variation within species, e.g., be the result of declining population sizes without actual extinctions. This should be detectable by losses of overall zOTU diversity and zOTU richness within single OTUs. To derive within OTU richness, we identified all OTUs that consisted of more than one zOTU. The richness of zOTUs within each of these OTUs was then calculated.

We also measured temporal abundance changes of single OTUs within sites. Within OTUs, temporal changes in read abundance at a site should reflect the relative abundance with reasonable accuracy ^61^. Only sites spanning a minimum time series of 10 years were included, and we only used species that occurred in at least three sampling events for a particular site and for which at least 100 reads were recovered. This filtering served to exclude rare species, which imitate abundance increases or declines by randomly occurring early or late in the time series. To account for likely fluctuations in abundance, we used the log+1 of read abundance. Significant increases or declines of abundance over time were estimated for each OTU and site using linear models (LM) in R.

Arthropods are an ecologically very heterogeneous taxon, with different groups showing very different life histories and possibly responses to ecosystem change. To account for this heterogeneity, we calculated *α*- and *β*-diversity metrics for the complete arthropod dataset, as well as the 10 most common arthropod orders in the dataset. Using NMDS (k = 2, 500 replications, Jaccard dissimilarity) in vegan, we visualized differentiation of the recovered arthropod communities. We then tested for effects of tree species, sampling year, sampling site, land use type, weather before sampling, amount of leaf material in a sample, climatic variables and detected pesticide load on *α*- and *β*-diversity. Also, hand-collected branch clipping samples, as well as the freeze-dried samples, were compared with the ESB samples for their recovered arthropod community composition and diversity.

Factors contributing to *β*-diversity were evaluated using a PERMANOVA in *vegan*. Statistical analysis for temporal changes of *α*-diversity and relative copy number were performed using the *lme4* (v 1.1-27.1; 2021) ^69^ package in R. Linear Mixed Models (LMM) were applied to analyze the statistical importance of involved fixed and random effects. Site ID and land use type were included as random effects. Temperature (annual, summer, and winter temperatures), corresponding rainfall data, year, and tree species were treated as fixed effects. The Akaike Information criterion (AIC) was used in stepwise regression to identify the final models. The final model for *α*-diversity only comprised the variables tree species and winter temperature (marginal R^2^ = 0.357, conditional R^2^ = 0.561, n = 312). Beech and pine were associated with the highest richness count. Despite a positive and significant statistical association between *α*-diversity and winter temperature (p = 0.022), the effect was small and mainly driven by the sample size: removing the tree species from the model and keeping only the winter temperature as fixed resulted in a marginal R^2^ of 0.013. The final model for estimating relative copy number resulted in a model comprising tree species and year as fixed factors (marginal R^2^ = 0.254, conditional R^2^ = 0.489, n = 312). The association with year was negative and highly significant (p < 0.001).

## Data availability

All raw reads are available in the Dryad Digital Repository (https://doi.org/10.5061/dryad.x0k6djhmp).

## Acknowledgments

We thank Karin Fischer for assistance with lab work. The German Environment Agency made the leaf samples available. Thanks to Diethard Tautz, Natalie Graham and Rosemary Gillespie for critical revision of an earlier version of the manuscript. Frank Thomas, Dorothee Krieger and Bernhard Backes provided access to the dried leaf homogenate we used here. Andrea Koerner helped in acquiring the leaf samples from the ESB.

## Author contributions

Conceptualization: HK

Methodology: HK, SW, RB, AM, SK, CM, SS, RW, RR, SK, JH

Data mining: RW, SW, HK, RR

Formal analysis: HK, TU, SW

Visualization: SW, SRK

Writing – original draft: HK, SW, SRK

Writing – review & editing: AM, RR, AH, CM, SS, JK, DT, RK, MP, MV

## Supplementary Information

### Supplementary Figures

**Fig. S1:**
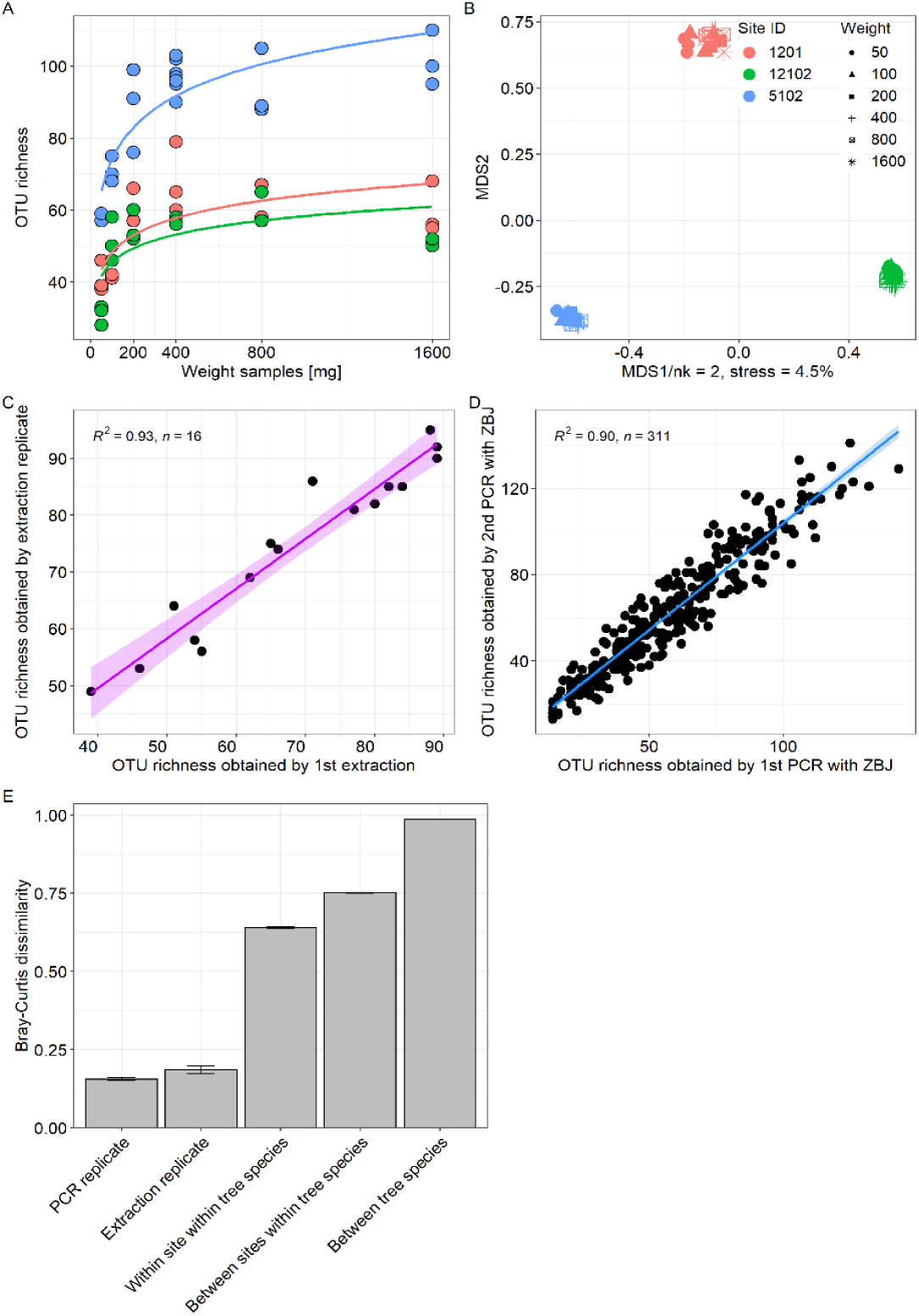
A) Saturation curves of recovered arthropod OTU richness for three sampling sites as a function of the amount of leaf homogenate used for DNA extraction. Extractions were made in triplicate for each input weight. B) NMDS plot for the same samples. Homogenate samples cluster by sampling site, while the amount of leaf homogenate used for DNA extraction has no discernable effect on β-diversity. C) Correlation of OTU richness obtained from extraction replicates. D) Correlation of OTU richness obtained from PCR replicates. E) Average β-diversity dissimilarity between PCR replicates, extraction replicates, different years within a site, different sites within a tree species and between tree species.

**Fig. S2:**
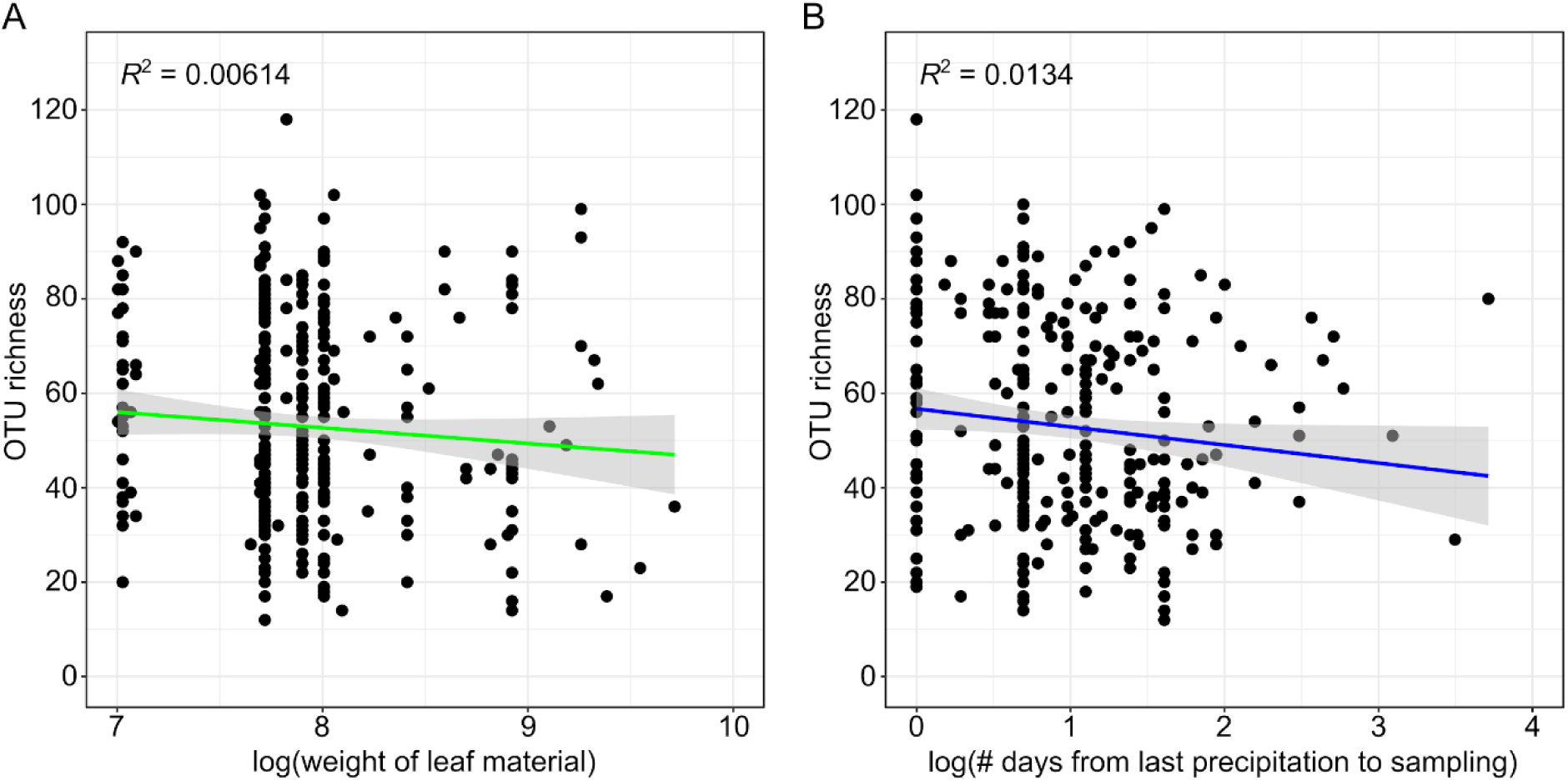
A) Correlation between the amount of leaf material (wet weight of leaves, in grams) in a homogenate sample and recovered arthropod community richness. No significant effect of increasing leaf amount on diversity was recovered, suggesting that the large number of leaves collected – several thousand per ESB sample – is sufficient to saturate the recovered diversity. B) Correlation between richness and the number of days between last precipitation event and sampling event. The number of days between rainfall and sampling did not affect the recovered diversity.

**Fig. S3:**
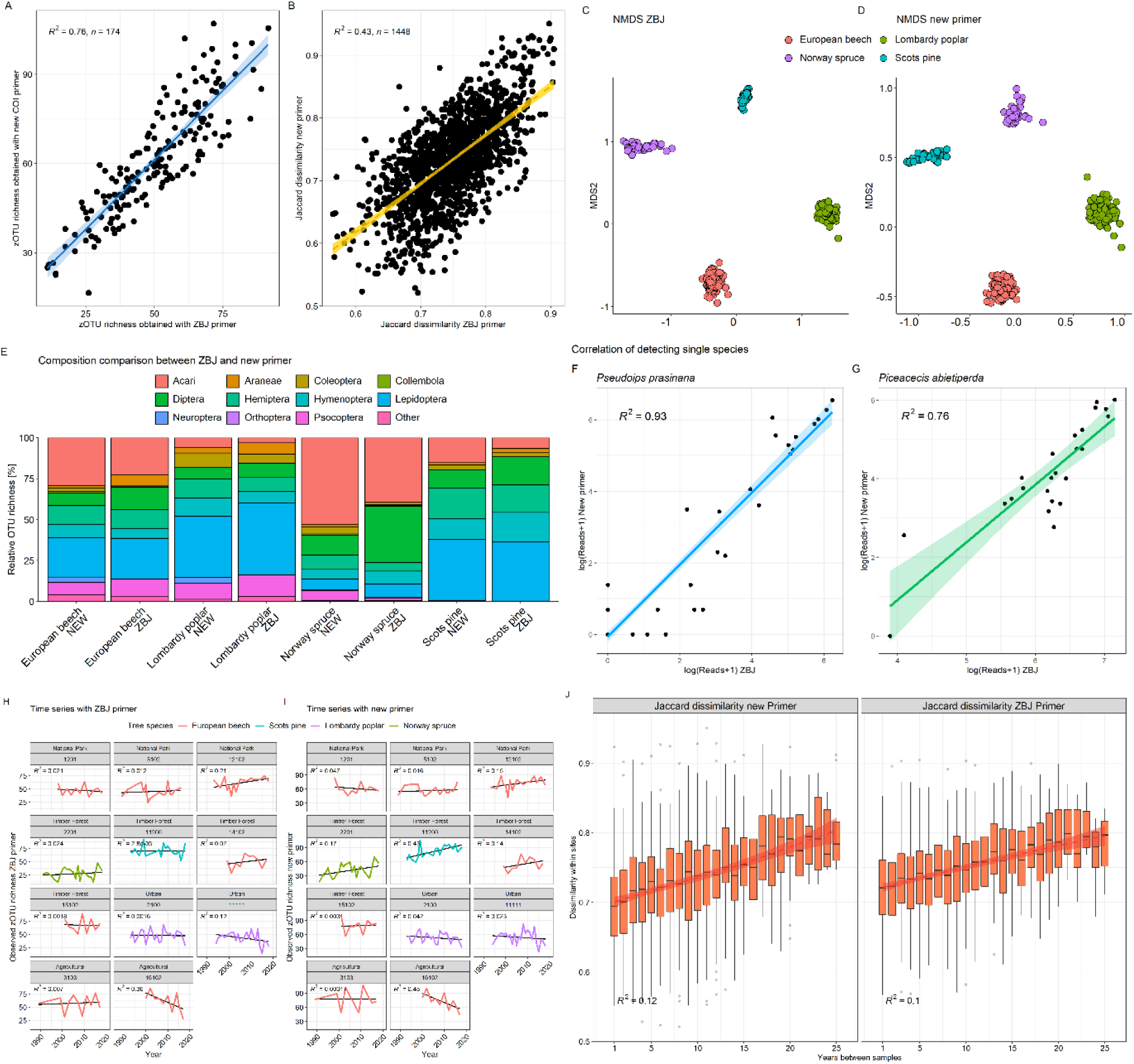
Comparison of recovered taxonomic composition and diversity patterns for the two COI markers (ZBJ-ArtF1c/ZBJ-ArtR2c vs. NoPlantF_270/mICOIintR_W) used in this study. A) Correlation of zOTU richness and B) β-diversity between samples. C-D) NMDS ordination of samples. E) Order-level taxonomic composition by tree species. F-G) Comparison of single species read abundances of two exemplary taxa. H-I) zOTU richness over time for the eleventime series analyzed for both markers. J) Temporal community turnover: Jaccard dissimilarity within sites compared against temporal distance between sampling events.

**Fig. S4:**
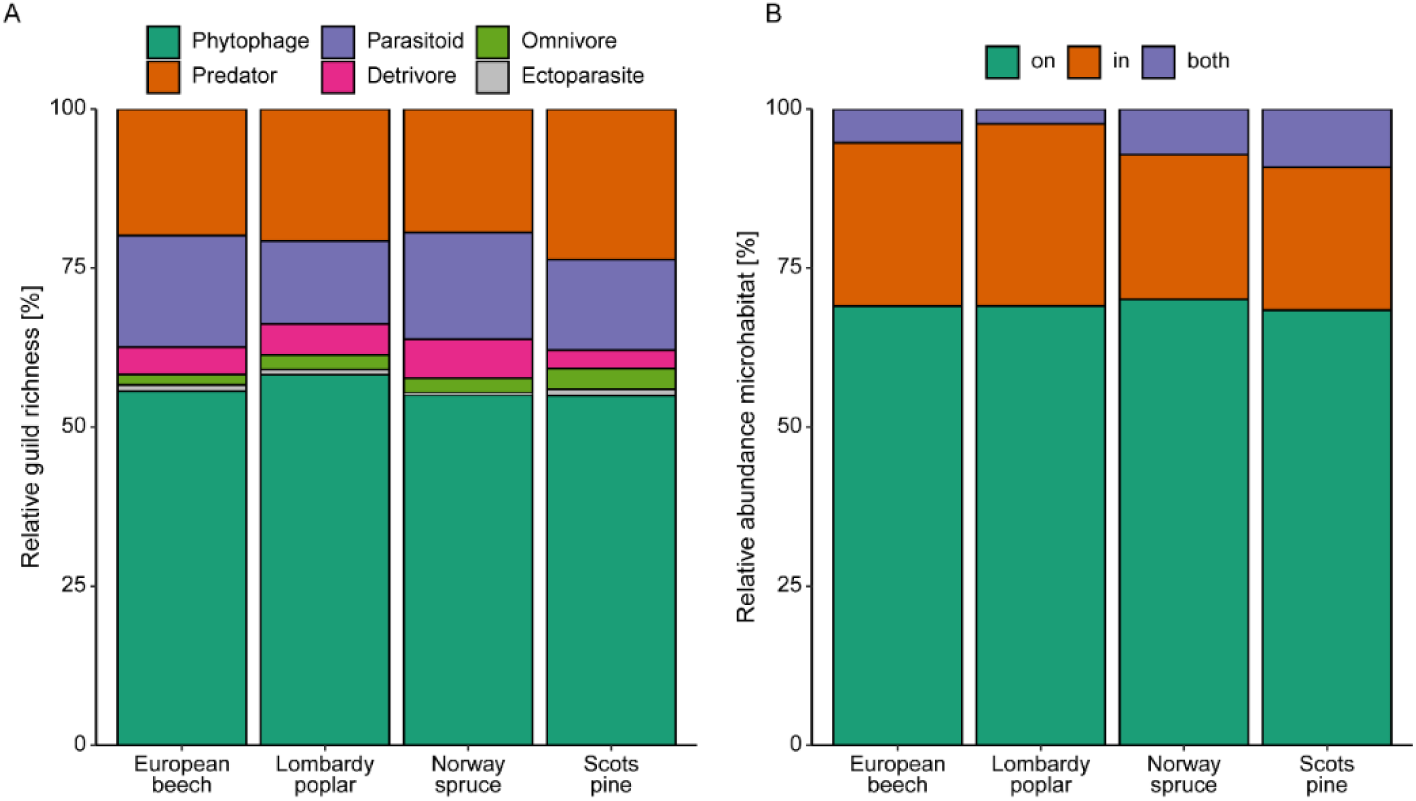
Ecological diversity of arthropods species recovered from the four tree species. A) Relative abundance of arthropod OTUs in different feeding guilds. B) Relative abundance of arthropod OTUs whose DNA likely originated from outside or inside the sampled leaves.

**Fig. S5:**
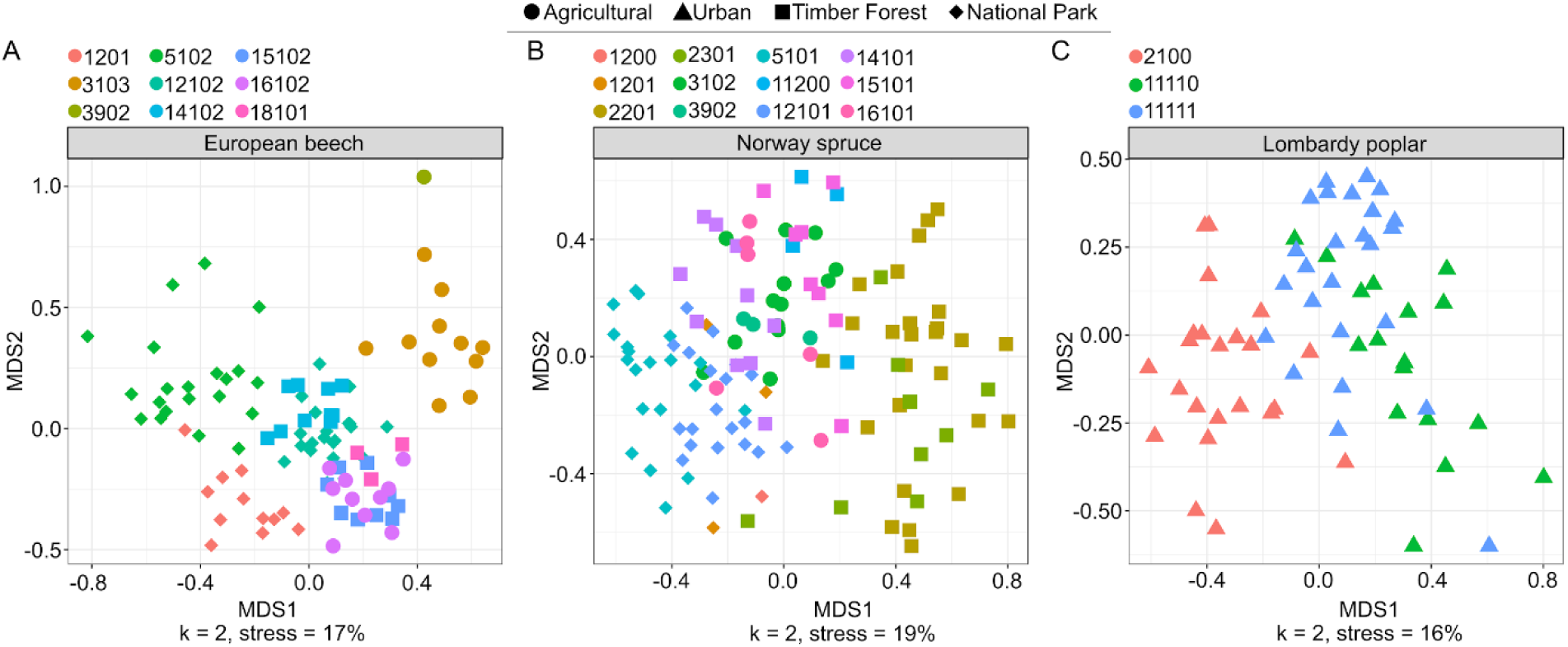
NMDS showing arthropod community differentiation by site, separated by tree species. A) European Beech, B) Norway Spruce, C) Lombardy Poplar. Color represents site and shape represents land use type.

**Fig. S6:**
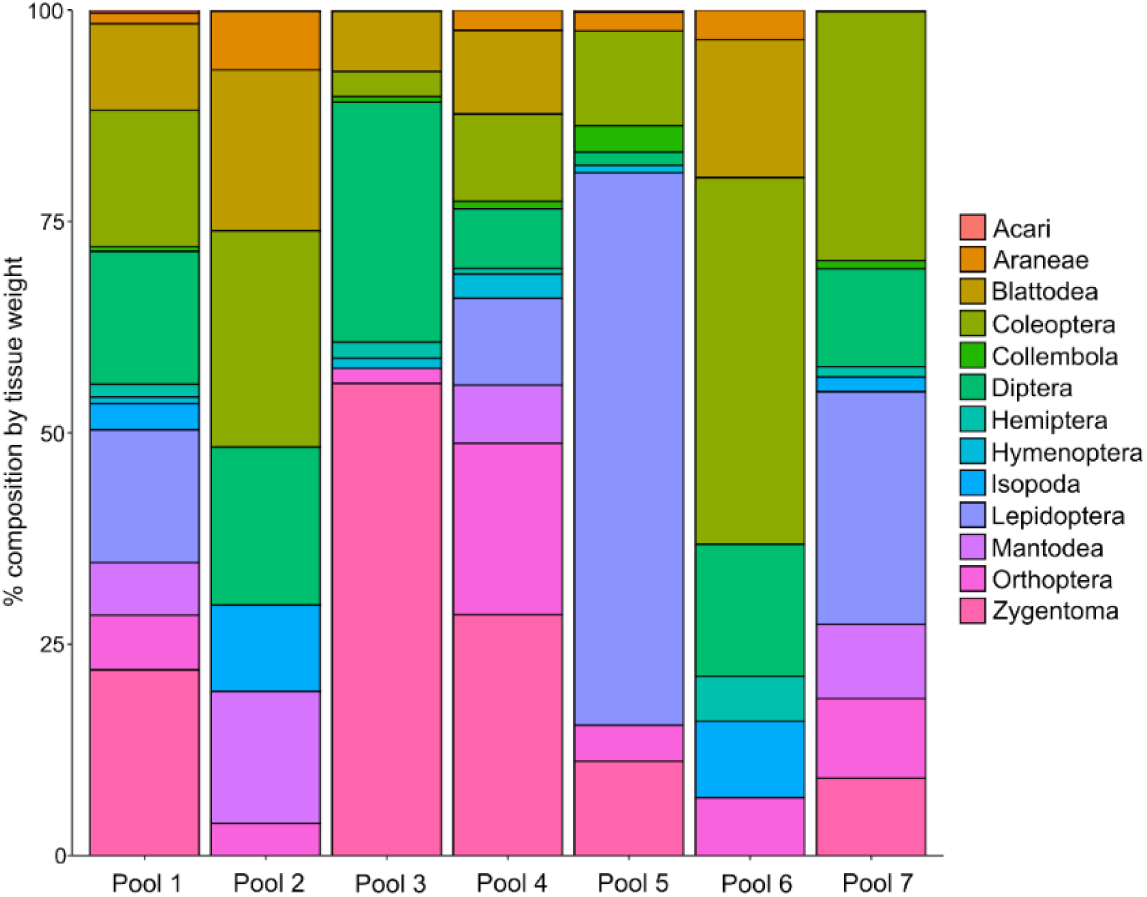
Order-level composition of the seven mock communities (Pools 1-7) used to test our qPCR assay. Each community contained different DNA proportions of species from 13 different orders and three classes across the arthropod tree of life.

**Fig. S7:**
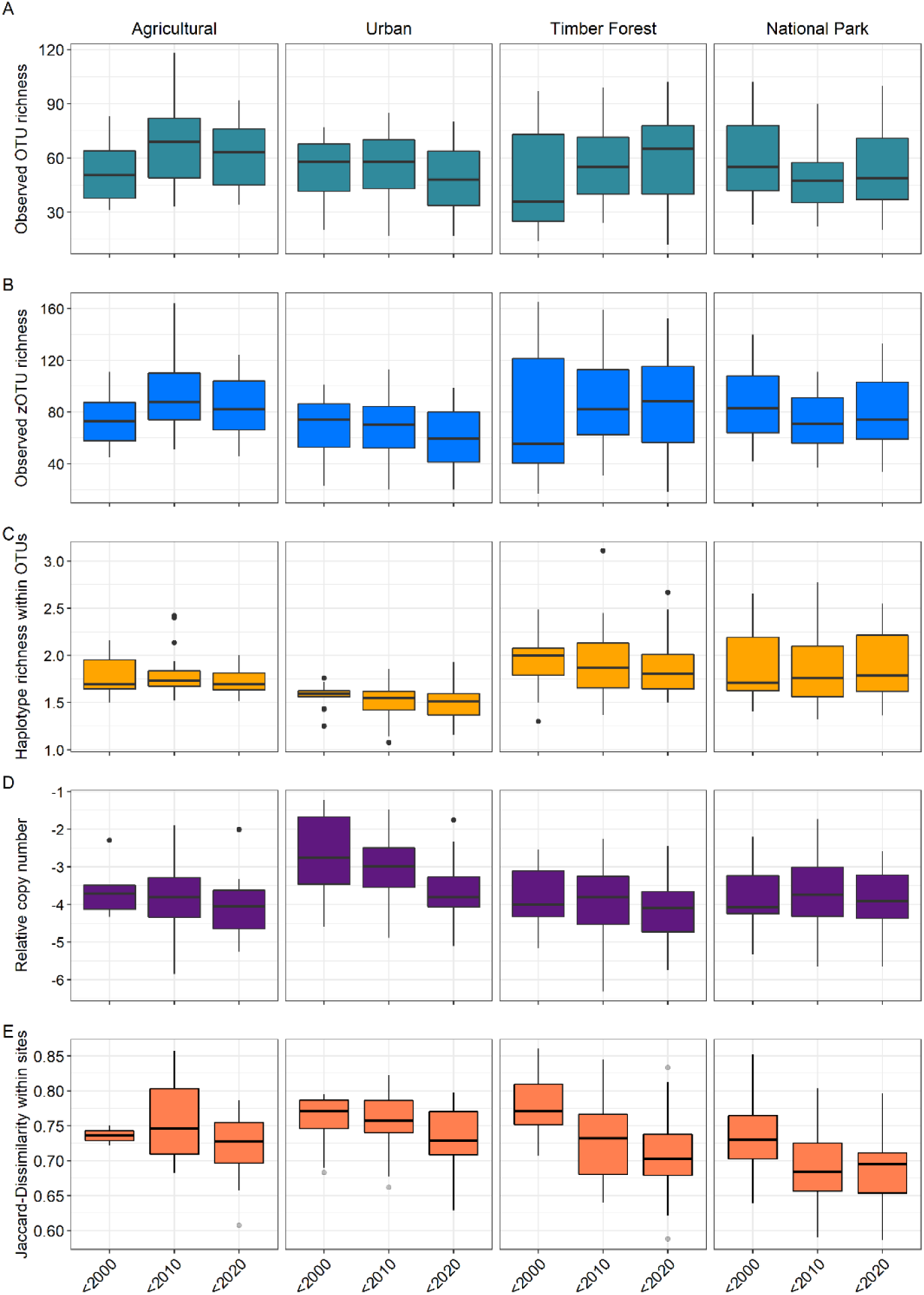
Metabarcoding based diversity indices and qPCR based relative copy number of arthropod rDNA per land use type over three decades. Boxplots of A) OTU richness, B) zOTU richness, C) haplotype richness within OTUs, D) relative copy number of arthropod rDNA, E) Jaccard dissimilarity between the given decade and the last sampling event in 2018, measured within sites. All datasets are merged by decade for clarity (before 2000, 2000 – 2009, and 2010 – 2018).

**Fig. S8:**
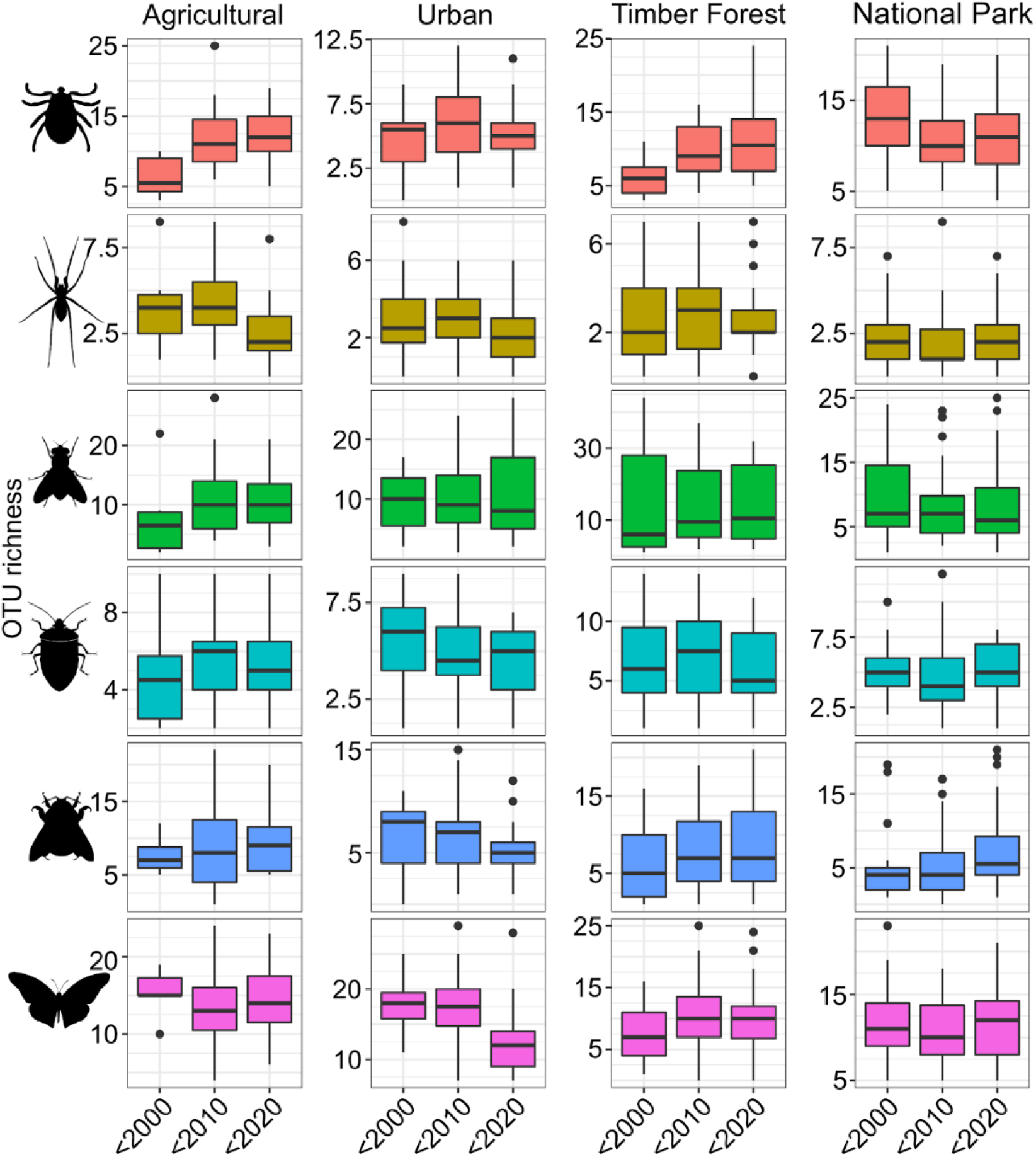
Boxplots of OTU richness by decade and land use type for the six most speciose arthropod orders in our dataset. The data are merged by decade for clarity (before 2000, 2000 – 2009, and 2010 – 2018). Richness is shown from top to bottom for Acari, Araneae, Diptera, Hemiptera, Hymenoptera and Lepidoptera.

**Fig. S9:**
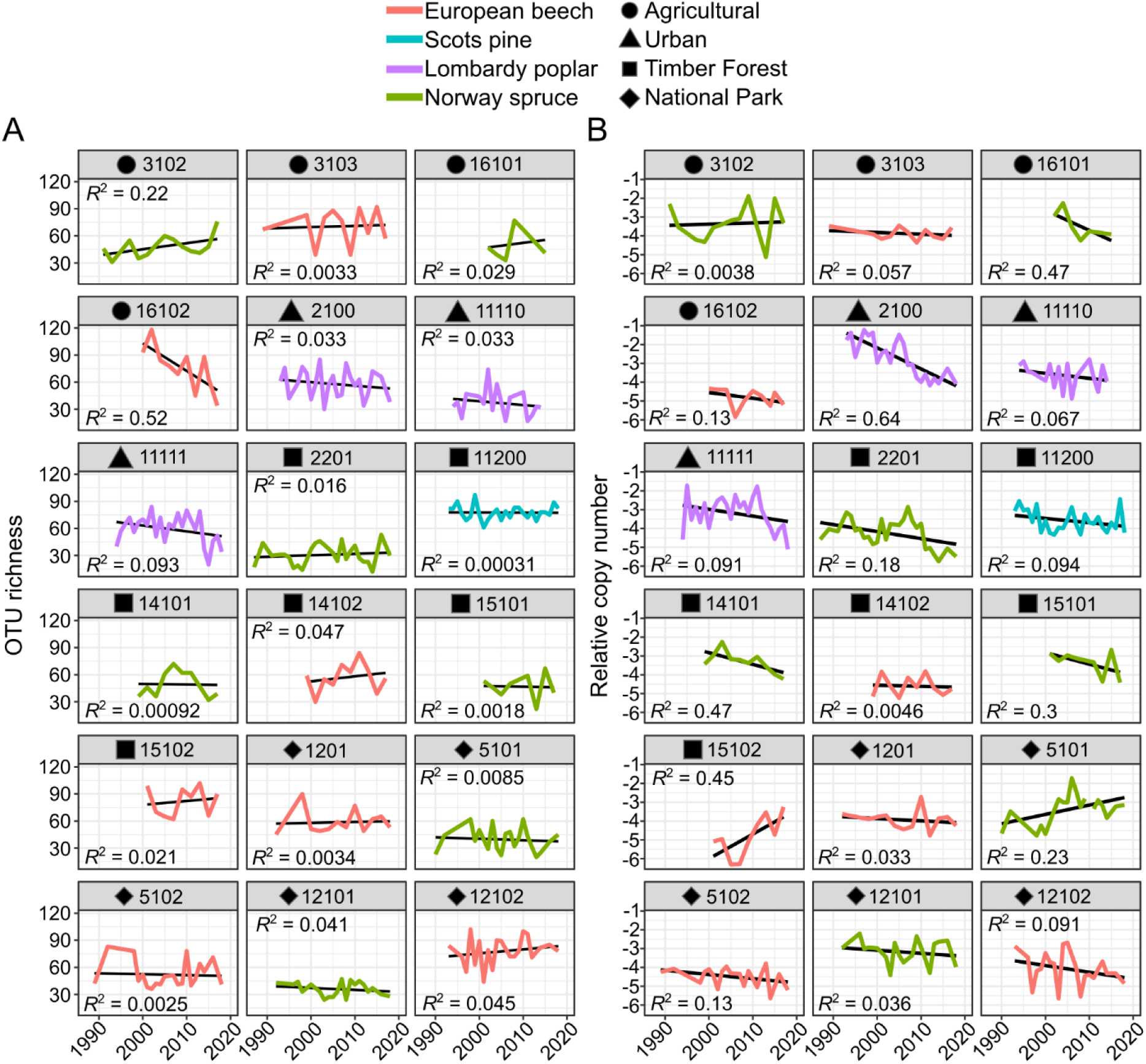
Arthropod diversity and relative copy number over time for all sites with time series longer than 10 years. A) OTU richness, B) Relative arthropod DNA copy number. Colors represent tree species; symbols represent land use types. Site name is indicated above each plot.

**Fig. S10:**
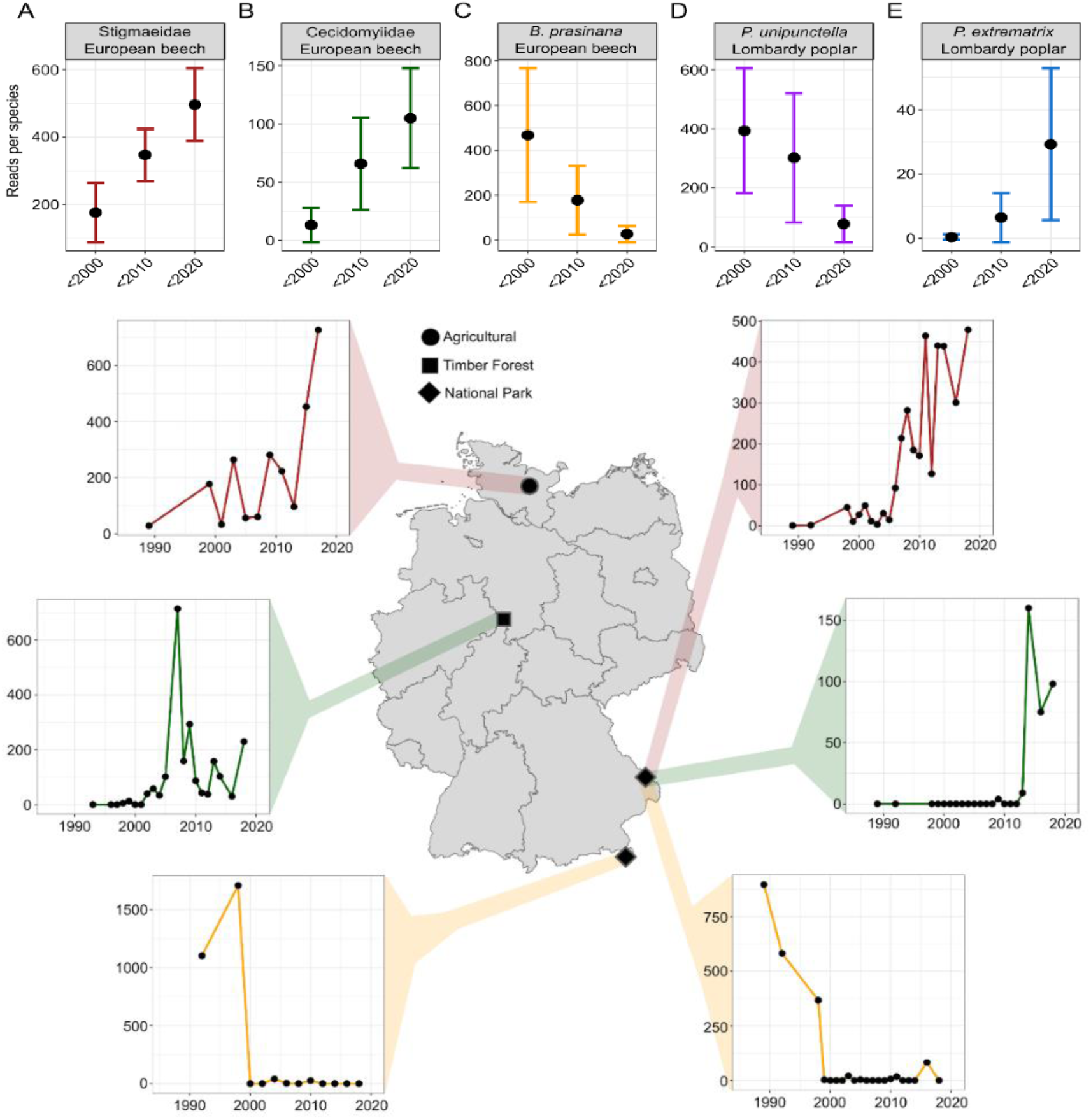
Temporal change in relative read abundance for five exemplary arthropod OTUs at all beech and poplar sites. A) Unidentified mite showing considerable increase over time. B) Unidentified gall midge also showing an increase. C) Decline in the Green Silver-Line over time. D) Decline in *Phyllocnistis unipunctella*, a leaf-mining lepidopteran, at poplar sites. E) Increase in a congeneric miner, *P. extrematrix*, at the same sites. F) Exemplary gains of the OTUs from A) and B) and losses of the OTU from C) at four sites in Germany.

**Suppl. Table 1:**
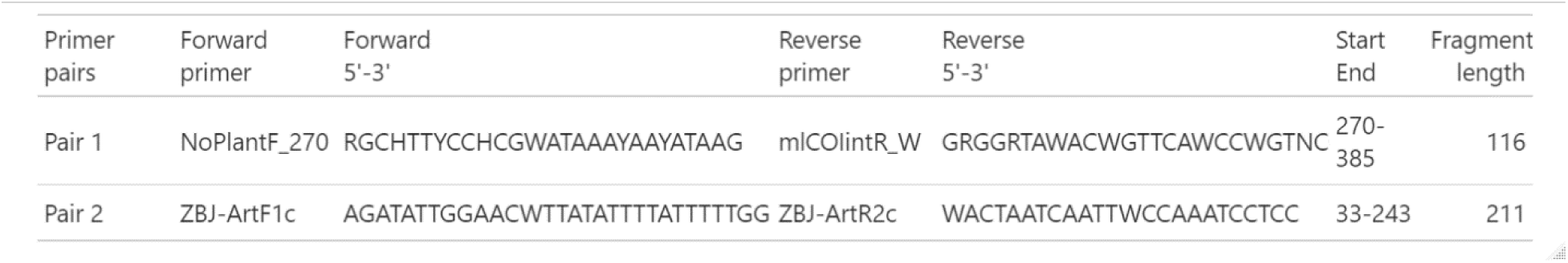
Primers used in this study. Start and end positions refer to the COI gene of the mitochondrial reference genome of *Drosophila melanogaster*.

Suppl. Table 2: OTU table with metadata

